# Diversity of dominant soil bacteria increases with warming velocity at the global scale

**DOI:** 10.1101/2020.04.27.063834

**Authors:** Yoshiaki Kanzaki, Kazuhiro Takemoto

**Author notes:** Corresponding author’s.

## Abstract

Understanding global soil bacterial diversity is important because of the key roles soil bacteria play in the global ecosystem. Given the effects of environmental changes (e.g., climate change and human effect) on the diversity of animals and plants, effects on soil bacterial diversity are expected; however, they have been poorly evaluated to date. Thus, in this study, we focused on the soil dominant bacteria because of their global importance and investigated the effects of warming velocity and human activities on their diversity. Using a global dataset of bacteria, we performed spatial analysis to evaluate the effects, while statistically controlling for the potential confounding effects of current climate and geographic parameters with global climate and geographic data. It was demonstrated that the diversity of the dominant soil bacteria was influenced globally by warming velocity (showing significant increases) in addition to aridity index (dryness) and pH. The effects of warming velocity were particularly significant in forests and grasslands. An effect from human activity was also observed, but it was secondary to warming velocity. These findings provide robust evidence, and advance our understanding of the effects of environmental changes (particularly global warming) on soil bacterial diversity at the global scale.

## Introduction

Understanding the global biogeography of microorganisms (particularly bacteria) in soils is a significant challenge as bacteria are present in all environments and soil bacteria play important roles in carbon and nutrient cycling, agriculture, animal (and also human) health, and food webs [1–3]; microbial diversity is particularly important in this context [3–6]. Previous work [1] investigating the global biogeography of soil bacteria found that only 2% of bacterial taxa account for almost half of the soil bacterial communities sampled globally. These dominant soil bacteria enhance our understanding of global soil bacterial diversity and distribution and are likely critical drivers, or indicators, of key soil processes worldwide [1].

Given the global importance of soil bacterial diversity, understanding associations between soil bacteria and their surrounding environments is also important and determining the effects of environmental changes (e.g., climate change and human effects) on soil biodiversity is of particular interest [2,7,8]. Historically, climate change (i.e., warming velocity) has been shown to decrease animal endemism [9], and human activities and their effects on the climate and environment are causing animal and plant extinctions globally [10–12], affecting ecosystem structures [13,14]. These results suggest that environmental changes will affect soil bacterial diversity. Several previous studies have reported these effects; however, microbes appear to show different responses to environmental changes to those of plants and animals. For example, observed positive associations between bacterial diversity and annual mean temperatures suggests that global warming increases diversity [15,16]; moreover, experimental climate warming increases grassland soil bacterial diversity [17] and accelerates its temporal scaling [18]. Human activities (e.g., land use) are known to positively affect bacterial diversity at local [19,20] and regional scales (e.g., across Europe [21]).

More focused investigations are required to understand the associations between environmental changes and bacterial diversity. Previous studies have mainly investigated continental-scale bacterial diversity, and evaluations of spatial autocorrelations between sites and variables have been understudied despite the known importance of these factors [13,14,22]. More importantly, the effects of environmental changes on bacterial diversity are still debatable because previous studies have focused on current temperatures. In particular, many evaluations have been based on the differences in current temperatures between sites rather than temperature changes over time within sites. This issue demonstrates the need for evaluating the effects of climate change and human activities at the global scale using spatial analysis. This is now possible because of the development of measurement technologies and improvement of infrastructures for databases that have increased the availability of global datasets of warming velocity [9] and human effects [23].

Controlling for potentially confounding effects of other environmental factors is also important. For example, it is well established that bacterial diversity is related to pH and dryness (e.g., measured using an aridity index calculated from mean annual precipitation and evapotranspiration) [1,3,15,24]. Precipitation seasonality [25], plant productivity [26,27], and potentially Ultraviolet (UV) radiation [28] all affect soil bacterial biodiversity. Climate change is known to influence these factors and global warming may increase all three [29–31]. Soil pH transitions from alkaline to acid when mean annual precipitation begins to exceed mean annual potential evapotranspiration [32]. Therefore, there is a need to break down the complex effects of different environmental factors on soil bacterial diversity. Although some previous studies have considered these confounding effects (e.g., [15]), limitations in environmental parameter data availability have resulted in a poor understanding generally.

In this study, we focused on dominant soil bacteria because of their global importance [1] and aimed to investigate whether environmental changes (i.e., warming velocity and human effects) affect soil bacterial diversity at the global scale. Using a global dataset of dominant bacterial phylotype compositions (constructed in [1]) and databases of environmental parameters, we comprehensively investigated how environmental changes contribute to bacterial diversity at the global scale while statistically controlling for potential confounding effects using spatial analysis.

## Material and Methods

### Dataset

We used a global dataset of the compositions of 511 dominant bacterial phylotypes (operational taxonomic units; OTUs) identified from soils collected from 237 locations across 6 continents [1] (see also figshare.com/s/82a2d3f5d38ace925492). To measure bacterial diversity, we computed richness, Simpson index, Shannon index, and species evenness at the OTU level using R statistical software (version 3.5.0; www.r-project.org) and the package *vegan* (version 2.5.6).

From the dataset in [1], we also extracted the aridity index (precipitation/evapotranspiration), mean diurnal temperature range (MDR; °C), net primary plant productivity (NPP; Normalized Difference Vegetation Index: NDVI [33]), precipitation seasonality (PSEA; coefficient of variation), pH, UV radiation index (unitless), ecosystem type (forest/grassland/shrubland), and latitude and longitude at each observation site. The dataset included 114 forest, 82 grassland, and 41 shrubland data that were analyzed as ecosystem specific datasets.

We obtained additional climate parameters based on the latitudes and longitudes of identified observation sites. In particular, the climate parameters were obtained based on the procedures established in our previous publications [13,14]. Annual mean temperature (AMT; °C) and temperature seasonality (TS; standard deviation) with a spatial resolution of 2.5’ were obtained from the WorldClim database [23] (version 2.0, release 1; www.worldclim.org). For evaluating human effects, we obtained the human footprint (HF) scores, which have a spatial resolution of 1 km grid cells, from the *Last of the Wild Project* [34] (version 3). The HF scores are defined based on human population density, human land use and infrastructure, and human access. Warming velocity (WV) also was computed. As in [9,35], we defined the velocity as the temporal AMT gradient divided by the spatial AMT gradient. The temporal gradient is defined as the difference between the current and past AMT. The past AMT was the CCSM3 model-based Last Glacial Maximum (LGM) AMT, available in the WorldClim database (www.worldclim.org/past). The spatial gradient was the local slope of the current climate surface at the observation site. We obtained the slope using the function *terrain* (with the option neighbors = 4) in the R package *raster* (version 2.9.5).

These data and parameters are available in the supporting information, S1 Dataset.

### Data analyses

The data analyses were based on the procedures described in our previous publications [13,14]. We performed regression analysis using R to evaluate the contribution of each variable to the diversity index. We considered both ordinary least-squares (OLS) regression and the spatial analysis approach (S1 Code).

For the OLS regression, we constructed full models encompassing all explanatory variables (AMT, Aridity index, HF, MDR, NPP, pH, PSEA, TS, UV, and WV), and selected the best model to obtain the most simplified model and to simultaneously avoid multicollinearity in the full model. The best model was selected based on the sample-size-corrected version of the Akaike information criterion (AICc) values using the R package *MuMIn* (version 1.43.6). When focusing on the best model only, however, the importance of certain variables may be overestimated; moreover, important variables may be overlooked. To avoid such a model selection bias, we adopted a model-averaging approach using *MuMIn*. We obtained the averaged model in the top 95% confidence set of models. A global Moran’s test was used to evaluate spatial autocorrelation in the regression residuals using the function *lm*.*morantest*.*exact* in the R package *spdep* (version 0.6.13).

As in [14,35,36], the richness and WV were log-transformed. Aridity index, PSAE, and NPP were also log-transformed for normality. The variables were normalized to the same scale, with a mean of 0 and standard deviation of 1, using the *scale* function in R before the analysis.

We also considered a spatial eigenvector mapping (SEVM) modeling approach [22,37] to remove spatial autocorrelation in the regression residuals. Specifically, we adopted the Moran eigenvector approach using the function *SpatialFiltering* in the R package *spatialreg* (version 1.1.5). As with the OLS regression analysis, we constructed full models, and then selected the best model based on AICc values. The spatial filter was fixed in the model-selection procedures [37]. We also obtained the averaged models.

The contribution (i.e., non-zero estimate) of each explanatory variable (i.e., environmental parameters) to bacterial diversity was considered significant when the associated *p*-value was less than 0.05.

We obtained the residuals of the explanatory variables and bacterial diversity according to the SEVM modeling approach-based best models.

## Results

To avoid redundancy, only the results of the Shannon index are presented in the main text (Table 1). The results of the other indices are available in the electronic supplementary material.

**Table 1.** Influence of explanatory variables on the diversity (Shannon index) of soil dominant bacteria.

| Variables | OLS |  |  | SEVM |  |  |
| --- | --- | --- | --- | --- | --- | --- |
|  | Estimate (Full) | Estimate (Best) | Estimate (Average) | Estimate (Full) | Estimate (Best) | Estimate (Average) |
| AMT | -0.028 (0.77) |  | -0.046 (0.47) | -0.136 (0.15) | -0.094 (0.09) | -0.120 (0.13) |
| Aridity index | -0.406 (<0.01) | -0.353 (<0.01) | -0.375 (<0.01) | -0.431 (<0.01) | -0.439 (<0.01) | -0.398 (<0.01) |
| HF | 0.051 (0.30) |  | 0.050 (0.29) | 0.103 (0.03) | 0.101 (0.03) | 0.098 (0.04) |
| MDR | 0.060 (0.42) |  | 0.021 (0.72) | 0.117 (0.11) | 0.094 (0.08) | 0.109 (0.07) |
| NPP | 0.058 (0.54) |  | 0.057 (0.48) | 0.147 (0.12) | 0.154 (0.05) | 0.148 (0.09) |
| pH | 0.430 (<0.01) | 0.429 (<0.01) | 0.429 (<0.01) | 0.481 (<0.01) | 0.467 (<0.01) | 0.466 (<0.01) |
| PSEA | 0.035 (0.60) |  | 0.005 (0.92) | 0.052 (0.45) |  | 0.035 (0.59) |
| TS | -0.071 (0.43) |  | -0.010 (0.88) | -0.083 (0.36) |  | -0.051 (0.59) |
| UV | -0.089 (0.40) |  | -0.049 (0.42) | 0.002 (0.98) |  | 0.034 (0.74) |
| WV | 0.292 (<0.01) | 0.303 (<0.01) | 0.295 (<0.01) | 0.273 (<0.01) | 0.233 (<0.01) | 0.261 (<0.01) |
| Moran's I | 0.148 (<0.01) | 0.162 (<0.01) |  | -0.088 (0.51) | -0.073 (0.49) |  |
| $R^2$ | 0.571 (<0.01) | 0.564 (<0.01) | | 0.661 (<0.01) | 0.658 (<0.01) | |

AMT, MDR, and TS indicate annual mean temperature, mean diurnal temperature range, and temperature seasonality, respectively; PSEA represents precipitation seasonality; NPP represents normalized difference vegetation index; UV represents UV radiation index; HF and WV represent the human footprint score and warming velocity, respectively. The estimates in the full, best, and averaged models based on the ordinary least squared (OLS) regression and spatial eigenvector mapping (SEVM) modeling approach are shown. $R^2$ denotes the coefficient of determination for the full and best models based on the OLS regression and SEVM modeling. Values in brackets are the associated $p$ -values.

The Shannon index was associated with several environmental changes (Table 1). Specifically, the Shannon index showed a positive correlation with warming velocity (Fig 1A) and HF score (human effects). The aridity index (i.e., dryness; Fig 1B) and pH (Fig 1C) were negatively and positively associated with the Shannon index, respectively. The aridity index, pH, and warming velocity were also associated with the other diversity indices (i.e., richness, Simpson index, and evenness); however, the contribution of human effects to these diversity indices was not statistically significant concluded (Tables A–C in S1 File). UV radiation and plant productivity were negatively and positively associated with evenness, respectively (Table C in S1 File) but showed no correlation with the other diversity indices (Table 1 and Tables A–C in S1 File). Annual mean temperature, mean diurnal temperature range, temperature seasonality, and precipitation seasonality showed no correlations with any diversity indices.

**Fig 1.**
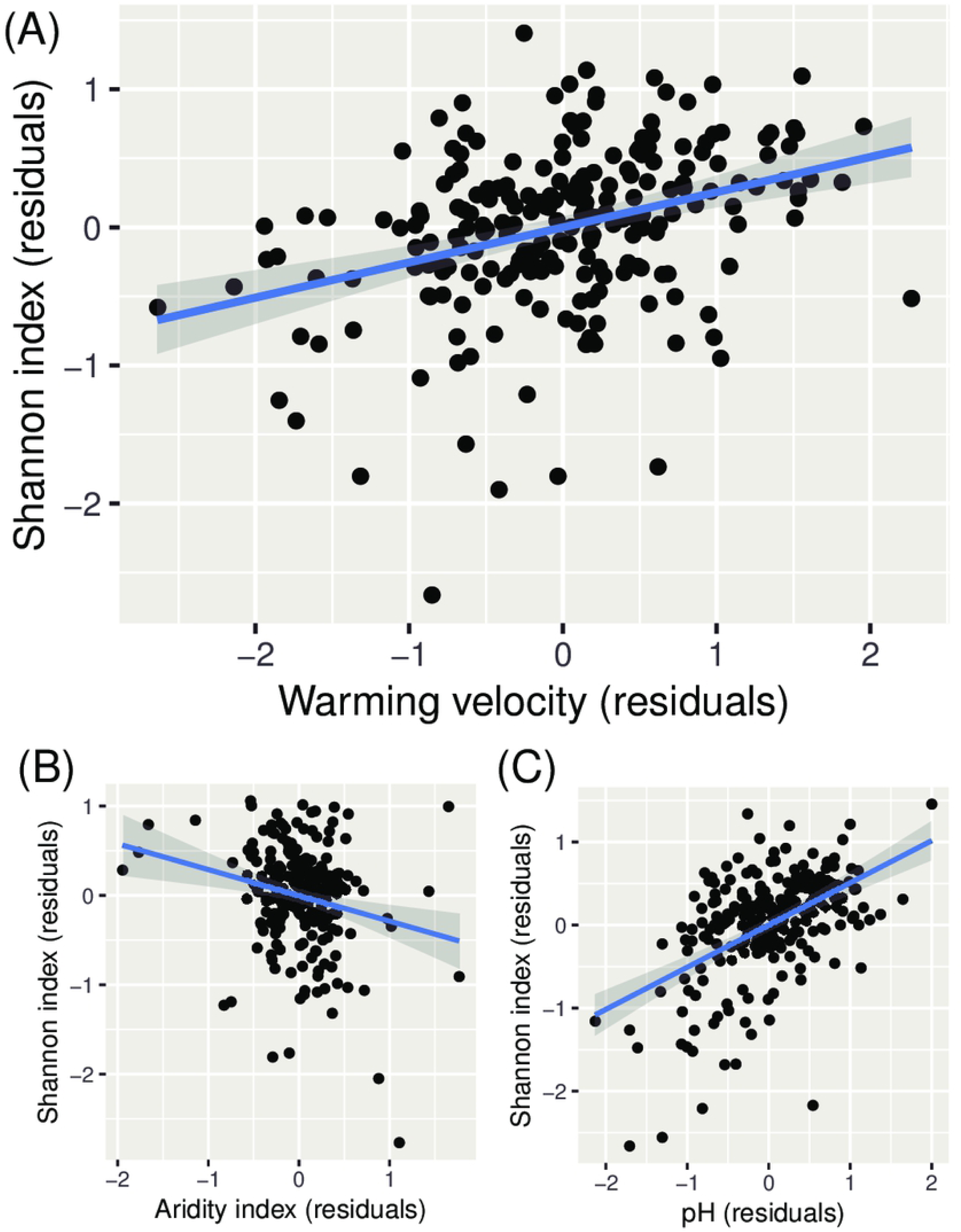
Scatter plots of the diversity (Shannon index) of soil dominant bacteria (residuals) versus environmental parameters (residuals). (A) Shannon index versus warming velocity. (B) Shannon index versus the aridity index. (C) Shannon index versus pH. The shaded grey area indicates the 95% confidence intervals around the regression lines.

In terms of the different ecosystem types, warming velocity was positively associated with all diversity indices in forests (Tables D–G in S1 File) and grasslands (Tables H–K in S1 File) but had no association with any diversity indices in shrublands (Tables L–O in S1 File). Human effects had a positive association with the Shannon index and richness in forests (Tables D and E in S1 File). A positive correlation between pH and bacterial diversity was observed without respect to ecosystem type (Tables D–O in S1 File). The aridity index was negatively correlated with all diversity indices in forests (Tables D–G in S1 File). Mean diurnal temperature range showed a negative association with evenness in shrublands (Table O in S1 File) and a positive association with the Simpson index and evenness in forests (Tables F and G in S1 File). Precipitation seasonality was positively associated with the Shannon index and richness in grasslands (Tables H and I in S1 File). Temperature seasonality showed a negative association with richness in forests (Table E in S1 File). Annual mean temperature, plant productivity, and UV radiation showed no correlation with any diversity indices when considering the ecosystem types (Tables D–O in S1 File).

## Discussion

Inspired by the attention given to the effects of environmental changes on soil biodiversity [2,7,8], we evaluated how environmental changes (warming velocity and human effects) influence the diversity of the dominant soil bacteria at the global scale while controlling for potential confounding effects using spatial analysis. Our analyses confirmed that environmental changes affect bacterial diversity. Specifically, we found that soil bacterial diversity increases with warming velocity and that this relationship is predominantly observed in forests and grasslands. This phenomenon was observed in multiple diversity indices; thus, we conclude that there is strong evidence for the observed relationship between warming velocity and soil bacterial diversity.

Our results are generally consistent with previous studies [15–18] that have concluded that warming results in increased soil bacterial diversity; however, our study provides complementary insights into these effects. Specifically, we emphasize the importance of warming velocity in this regard [9]. Previous studies [15,16] have primarily shown the effects of annual mean temperature; however, our analysis (Table 1) did not conclude any association between annual mean temperature and soil bacterial diversity. This discrepancy might be due to differences in the datasets and methods for data analyses between our work and previous studies. Our focus on global-scale bacterial diversity and the incorporation of control for the effects of multiple environmental factors addressed the aforementioned limitations of previous work (see Introduction). When preforming simple pairwise correlation analysis, a positive correlation between soil bacterial diversity (Shannon index) and annual mean temperature was also observed (Spearman’s rank correlation coefficient *r*_*s*_ = 0.26, *p* = 5.8e–05) in our dataset, which is consistent with the previous studies. Therefore, our analysis calls into question the previously observed association between annual mean temperature and soil bacterial diversity and instead highlights the importance of warming velocity along with other environmental factors.

Previous studies [17,18] reporting soil bacterial diversity increases with experimental climate warming have focused on local scales. Our results indicate that the positive effects of warming on soil bacterial diversity occur at the global scale and are obtained from real-word data. However, the effects are limited to data from historical climate change (warming velocity) while the focus in [17,18] was on current human-driven climate change. In this study, the warming velocity was considered to investigate the effect of climate change because global data on current climate change were unavailable. As described in our previous publications [13,14], the time-scale of the warming velocity may be overly long, compared to that of microbial community assemblages because the velocity was estimated based on the difference between the current and past (LGM) annual mean temperature. However, strong climatic shifts that have occurred since the LGM (21,000 BP) has affected geographical patterns of species endemism [9]. This indicates that species (and also bacterial) compositions are more sensitive to environmental changes in areas that have experienced these climatic shifts. In fact, factors driving shifts in soil bacterial communities can reflect the historic climate because soil properties change slowly over time [38].

Positive associations between warming and soil bacterial diversity are often explained by the metabolic theory of ecology [39,40]. Metabolic rate (oxygen consumption and energy demand) is an important physiological parameter in (microbial) ecology [41] because it can be used to estimate, and therefore understand, energy metabolism [42], population growth rate [43], genetic mutation rate [44,45], and species diversity [46,47]. Importantly, increases in temperature accelerates the metabolic rate [48], although this effect might reach a saturation point [49,50]. Thus, the theory predicts that climate warming will increase the rates of ecological and evolutionary processes [15,18], including the rates of genetic mutation, speciation, and interactions. In fact, community-level respiration of prokaryotic microbes may rise with global warming [51]. However, our results (Table 1) indicate that soil bacterial diversity is influenced by warming velocity rather than only by environmental temperature, implying that increases in soil bacterial diversity will not occur at low warming velocities despite high environmental temperatures. We believe that the theory can explain the positive association between warming velocity and soil bacterial diversity because the velocity is based on the difference between current and past annual mean temperatures. However, the theory (specifically, the equation used in [15]) may need to be modified to consider warming velocity.

Our results (Table 1) also indicate that human effects can positively affect soil bacterial diversity at the global scale, corroborating findings at local [19,20] and regional scales [21]. Human movement may facilitate the invasion of diverse (soil) bacteria from one area to another, and increases in habitat patches driven by human land use may increase soil bacterial diversity. It has been shown that soil bacterial diversity is positively associated with human population [52]. However, the direct effects of human effects are currently ambiguous because they depend on the types of diversity indices and ecosystem types investigated (see section 3). The effects of the other environmental factors (UV radiation, plant productivity, temperature seasonality, and precipitation seasonality) at the global scale may also still be uncertain because they were influenced by the types of diversity indices and ecosystem types included in the analyses.

Dryness (based on the aridity index results) and pH appear to affect soil bacterial diversity. Our findings that aridity index and pH had negative and positive correlations with bacterial diversity, respectively, are consistent with a number of previous studies [1,3,15,24], despite the limitations in terms of data and statistical analysis in these works (see section 1). This suggests that the evidence for these effects are robust.

The present analysis has several limitations. For example, our data are limited to the dominant soil bacteria, which we focused on because of their global importance [1]. However, this limitation may not pose a significant problem because similar tendencies between dominant bacterial taxa and all taxa generally have been observed [1] (e.g., a diversity index of all taxa was strongly correlated with that of the dominant taxa). Additionally, our study did not evaluate the relationships between specific phylotypes (e.g., genus and class) and environmental factors. This was because the main aim of this study was to investigate bacterial diversity; however, it was also owing to the difficulty in data analysis. The bacterial compositional data were high dimensional (511) compared to the sample size (235), and statistical analyses may not be applicable to compositional data generally because the assumption of independent variables may not be satisfied owing to the constant sum constraint [53,54]. More importantly, spatial autocorrelation also needs to be considered. No appropriate method for high-dimensional, spatial, and compositional data was available. To avoid these limitations, we simply focused on diversity indices, a compressed expression of bacterial compositions.

In addition, our results are limited by available datasets, as are many other geographic studies on soil bacteria. Global-scale datasets are now available owing to remarkable development in high-throughput sequencing techniques; however, the current amount of available data may still be insufficient to produce a definitive picture. Significant gaps in soil biodiversity data remain across northern latitudes, including most of Russia and Canada [55] and data are lacking from much of central Asia and central Africa, as well as many tropical regions. To account for these limitations, a more global-scale dataset on soil bacteria and environmental parameters should be constructed in the future. In this context, the Earth Microbiome Project [56] is significant, and data sharing generally [57] will be particularly important for future studies.

In conclusion, despite the limitations in our data analyses, our findings enhances our understanding of the effects of environmental changes (particularly global warming) on soil bacterial diversity.

## Acknowledgements

The authors would like to thank Editage (www.editage.com) for English language editing.

## Supporting Information

**S1 Dataset. Data on the diversity of the dominant soil bacteria and environmental parameters that were used for the analyses in this study**. The following table lists the Shannon index, Simpson index, richness, evenness, pH, Aridity index, annual mean temperature (AMT), mean diurnal temperature range (MDR), temperature seasonality (TS), precipitation seasonality (PSEA), UV radiation index (UV), normalized difference vegetation index (NPP), human footprint score (HF), warming velocity (WV), continent, and latitude and longitude of the observation site. See also the original dataset (figshare.com/s/82a2d3f5d38ace925492) for further details.

**S1 Code. R code used in our data analyses**.

**S1 File. Supporting Tables**. Tables A–O are included.

